# Neural signatures of second language proficiency in narrative processing

**DOI:** 10.1101/2022.10.14.512249

**Authors:** Ruiqing Zhang, Jing Wang, Hui Lin, Nicholas B. Turk-Browne, Qing Cai

## Abstract

Making sense of speech in a second language relies on multiple abilities. Differences in brain activity related to proficiency in language tasks have often been attributed to processing demands. However, during naturalistic narrative comprehension, listeners at different proficiency levels may form different representations of the same speech. We hypothesized that the synchronization of these representations across people could thus be used to measure second-language proficiency. Using a searchlight shared response model, we found that highly proficient participants showed synchronization in regions similar to those of native speakers, including in the default mode network and in the lateral prefrontal cortex. In contrast, participants with low proficiency showed more synchronization in auditory cortex and word-level semantic processing areas in the temporal lobe. Moderate proficiency showed the greatest neural diversity, suggesting lower consistency in the source of this partial proficiency. Based on these synchronization differences, we were able to reliably classify the proficiency level or predict behavioral performance on an independent English test in held-out participants, suggesting the identified neural systems represented proficiency-sensitive information that was generalizable to other individuals. These findings suggest higher second-language proficiency leads to a more native-like neural processing of naturalistic language, including in systems beyond the cognitive control network or the core language network.

**Highlights:**

- Neural synchronization in second-language speech processing reflects proficiency.
- High-proficiency individuals neurally resemble native speakers.
- Low-proficiency individuals are synchronized in perceptual and word semantics areas.
- Proficiency level can be predicted using neural synchronization signatures.

## Introduction

We rely on shared knowledge to understand each other, yet knowledge sharing relies on effective communication. A story “resonates with us” not only metaphorically but also physically: During perception or recall of continuous complex narratives, multiple neural systems track the transitions of events at different timescales (Baldassano et al., 2018, 2017). Neural responses to the same narrative are synchronized across multiple audience or readers (Lerner et al., 2011; Regev et al., 2019; Simony et al., 2016) and across storytellers (Silbert et al., 2014). Higher speaker-listener neural coupling predicts successful communication (Stephens et al., 2010). Shared knowledge has also been associated with synchronized event-specific response profiles across people recalling similar events (Chen et al., 2017), between encoding and recall processes (Zadbood et al., 2017), and between movie watching and speech perception (Nguyen et al., 2019; Zadbood et al., 2017). Different scales of information are represented in different neural substrates (Lerner et al., 2011) and at different orders of neural dynamics (Owen et al., 2021).

Various factors affect cross-individual synchronicity, such as attention (Regev et al., 2019) and interpretation of the materials (Nguyen et al., 2019). In the processing of speech in a second language (L2), these factors may reflect stable individual differences. When people listen to a talk in a non-native language, language proficiency affects the degree to which they can track the speech stream, follow the topic, and grasp various details and implications. According to Common European Framework of Reference (CEFR; www.coe.int/en/web/common-european-framework-reference-languages), basic users can understand phrases related to areas of their immediate priority if the articulation is clear and slow, whereas proficient users can understand extended discourses on complex and less familiar topics with only occasional missing details.

Three lines of findings on L2 or bilingual processing are most relevant to the present study. First, consistent with the convergence hypothesis (Green, 2003), L1 and L2 processing share cognitive mechanisms and neural substrates for speech perception, semantic representation, and syntactic processing (Abutalebi, 2008; Costa and Sebastian-Galles, 2014; Illes et al., 1999; Kotz, 2009; Perani et al., 1998; Sebastian et al., 2011). Second, L2 proficiency or ability has been associated with cognitive and neural activational differences that cannot be accounted for by the age of acquisition (Chee et al., 2001; Hamrick, 2015; Hamrick et al., 2018; Perani and Abutalebi, 2005; Stein et al., 2009; Wartenburger et al., 2003). Third, these neural functional differences between the use of languages are often attributed to the difference in processing demand: greater efforts are required for L2 or bilingual processing, either due to the less frequent use of the language, the transfer between linguistic knowledge, or the involvement of additional cognitive control/monitoring compared to monolinguals (Abutalebi, 2008; Berken et al., 2016; Costa and Sebastian-Galles, 2014; Golestani et al., 2006; Grant et al., 2015; Green, 2003; Luk et al., 2012; Weber et al., 2016).

Here we further propose that during naturalistic narrative processing, variation in language proficiency leads to not only cognitive control differences, but also diverse understanding and representations of the same speech. These diversified representations will disrupt the neural synchronizations across individuals, especially in brain areas associated with narrative processing. We hypothesize that people with a similar proficiency level will have similar neural response patterns, which differ from those at a different proficiency level. Lower proficient individuals may be synchronized in earlier perceptual and semantic areas such as auditory cortex and ventral temporal cortex, whereas higher proficiency individuals may be synchronized in later regions involved in narrative comprehension such as lateral and medial prefrontal cortex, temporoparietal cortex, and posterior cingulate cortex/precuneus (Chen et al., 2017; Honey et al., 2012; Lerner et al., 2011; Nguyen et al., 2019; Simony et al., 2016; Zadbood et al., 2017). Additional candidate areas include the posterior middle frontal gyrus and the precentral-postcentral areas (Honey et al., 2012).

The present study uses shared response model (SRM; Chen et al., 2015; Zhang et al., 2016) to investigate the neural signatures of second language proficiency during narrative comprehension. Complex naturalistic speech materials, namely TED talks, were used to examine higher-order processes that may not be easily observed in language tasks that tap into segregated cognitive variables. Identifying cross-individual synchronization in speech processing can be challenging because fine-grained semantic encoding patterns in the brain bear considerable individual differences (e.g., Huth et al., 2016). We apply SRM to partition multi-subject fMRI data into the common neural response profiles and subject-specific local functional topographies. The approach of using SRM in a searchlight has been validated by showing optimal performance in extracting consistent neural profiles within a group of individuals (Chen et al., 2016).

## Materials and methods

### Participants

A total of 47 healthy participants (32 females, mean age = 22.7 years, SD = 2.7) were recruited from the East China Normal University community. Participants were native Mandarin Chinese speakers with English as the second language (mean age of acquisition in listening = 8.7 years, SD = 2.7), right-handed, reported normal hearing and had no history of neurological disease or language impairment. Language history was collected using Language History Questionnaire version 2.0 (Li et al., 2014). All participants provided informed consent that was approved by the East China Normal University Institutional Review Board.

### Stimuli and experimental design

Stimuli of the main study were the audio tracks of 4 TED talks (www.ted.com), the topics of which were communication skills (“Talk nerdy to me” by Melissa Marshall; “Try something new for 30 days” by Matt Cutts; “Want to be more creative? Go for a walk” by Marily Oppezzo; and “Keep your goals to yourself” by Derek Sivers). The speeches lasted for 4.25 min (averaged speed 170 words/min), 6.48 min (60 words/min), 3.17 min (80 words/min) and 2.98 min (82 words/min), respectively. Participants were asked to listen to the talk with eyes open while being scanned. The four talks were presented in four separate scan runs, synchronized with MRI acquisition onset using PsychoPy (Peirce, 2007). To ensure that the participants paid attention to the stimuli, participants were asked to try their best to verbally report the content of the preceding talk in their native language after each run.

### Behavioral assessment of second language proficiency

Participants performed two behavioral tests before the main experiment. Phonological perception ability was measured using a phoneme discrimination task. In each trial, participants heard two word-like sounds in sequence and judged whether the two items sounded the same. The sounds were English words in the form of consonant(s)-vowel-consonant(s) pronounced by a native English speaker. Each pair was either the same or differed by one phoneme.

Language proficiency was estimated using the Cambridge assessment of English on listening (https://www.cambridgeenglish.org/exams-and-tests/). We used the materials that corresponded to the CEFR level A2, B1, B2, and C1. Participants listened to different paragraphs and answered questions about the materials. Performance was used to assign participants into the high, moderate, and low proficiency groups, as described in the *Results* section.

### MRI acquisition

Whole-brain images were collected using a 3.0-Tesla Siemens Trio MRI scanner (Siemens, Erlangen, Germany) with a 64-channel head coil. Functional images were acquired with an interleaved multiband echo-planar imaging sequence (TR = 1000 ms, TE= 32.6 ms, FoV = 192 mm, 2 mm isotropic voxel, 55° flip angle, 72 transversal slices, multiband factor 6). For offline correction for the susceptibility-induced distortions on EPI images, a pair of spin echo volumes was acquired in opposing phase encode directions (anterior to posterior and posterior to anterior) with EPI-matching slice prescription, voxel size, field of view, bandwidth, and echo spacing. Pulse and respiration during task were collected using a pulse detector on a finger and a respiration detection belt wrapped around the chest. T1-weighted anatomical image was obtained using a MPRAGE sequence (TR = 2530 ms, TE = 2.34 ms, FoV = 256 mm, 7° flip angle, 192 slices). Participants wore MRI-compatible passive noise-canceling headphones.

### Data analysis

#### Preprocessing

Images were preprocessed using FSL FEAT 5 (https://fsl.fmrib.ox.ac.uk/fsl/fslwiki/FEAT). Susceptibility-induced distortions were corrected using FSL TOPUP (Andersson et al., 2003). Off-resonance field image was derived from the two spin-echo field mapping images, with unit converted to angular frequency, and then submitted for B0 unwarping. Functional images were spatially realigned, distortion corrected, registered to the structural image and normalized to MNI152 template, temporally filtered (128 sec high-pass filter), and smoothed with a 5mm FWHM Gaussian kernel. Respiration and pulse recordings were processed using FSL PNM (https://fsl.fmrib.ox.ac.uk/fsl/fslwiki/PNM) to generate the covariates of physiological noise, which were regressed out from the preprocessed signal along with the head motion parameters.

#### Searchlight SRM

The subsequent analyses were performed using customized python scripts and the Brain Imaging Analysis Kit (Kumar et al., 2022; http://brainiak.org). In general, the SRM technique projects multi-subject data into a common, low-dimensional feature space. Given time-by-voxel matrices, SRM finds a voxel-by-feature transformation matrix for each subject and a feature-by-time matrix that is shared across all subjects. For this study, to identify the neural signatures for different levels of proficiency, shared response modeling was applied in a leave-one-participant-out cross-validation procedure within each searchlight cube. The cube contained 27 voxels and moved by a step of one voxel. The number of shared neural features was set to 10 (Chen et al., 2015). In each cross-validation fold in each searchlight location, three SRMs, one per proficiency group, were separately trained using all but the left-out participants’ data within each proficiency group. Three shared neural feature matrices (feature × time point) were learned. The feature matrices were applied separately to the left-out participant to reconstruct three sets of neural responses over time. These reconstructed responses represented three different predictions of the held-out participant’s neural response based on three different hypothesized proficiency levels. The reconstructed signals were averaged across all the voxels within the searchlight cube. Three Pearson’s correlation scores were computed between the original signal and each of the mean reconstructed time series. If the reconstruction derived from the actual proficiency group of the participant presented the highest correlation, this searchlight location was considered to be proficiency-sensitive.

For each participant, the correlation scores of these proficiency-sensitive locations were transformed to Fisher’ Z scores. The normalized correlation maps were averaged within each proficiency group. The maps were further thresholded against a null distribution generated by a 1000-iteration random permutation (Winkler et al., 2014). The resulting maps indicated the voxels that were reliably identified as being proficiency-sensitive across participants at the same proficiency level.

To further identify regions that were exclusively characteristic to one proficiency group, the pairwise differences between the mean Z maps were computed between each pair of proficiency groups. These difference Z maps were thresholded against a 1000-iteration random permutation. The thresholded maps showed voxels of which the response profiles were more characteristic for one proficiency level than the other.

#### Classification of proficiency level

To verify the sensitivity to proficiency of the brain maps revealed by searchlight SRM, classifiers were trained on the correlation maps to identify the proficiency level of a participant. In each cross-validation fold, one participant was left out. The training exemplars were the correlation between the actual data of each participant in the training set and the time series reconstructed based on the shared response of this training participant’s proficiency group. Support vector machine classifiers were trained and applied to the three correlation maps of the left-out participant to identify the group membership. A binary absolute accuracy score was determined by whether the predicted group label matched the actual, behaviorally determined label of the participant. The theoretical chance-level accuracy was 0.33. To statistically test whether the mean accuracy over participants were above chance level, the classification procedures were repeated on randomly shuffled labels of the language proficiency for 100,000 times to form a null distribution of the classification accuracy.

#### Prediction of second-language comprehension scores

The machine-learning procedure was the same as the classification analysis, except that a ridge regression model was trained to estimate the relation between the correlation patterns and the English comprehension scores of the training participants. The trained model was applied to correlation pattern of the left-out participant to predict the comprehension score. Pearson’s correlation between the predicted scores from all the cross-validation folds and the actual scores over participants was calculated as the metric of model performance.

## Results

### Behavioral results and assignment of proficiency groups

Participants were assigned to either low proficiency level (LP), moderate proficiency level (MP), or high proficiency level (HP) group based on their Cambridge listening comprehension test scores. The mean score on Cambridge listening comprehension test over participants was 4.27 out of 6.5 (65.38%), SD = 1.38. Participants were assigned to LP if the score was 1 SD below the mean or lower and to HP if their score was 1 SD above the mean or higher, and to MP otherwise (Figure 1). This resulted in 15 participants in LP (scores ranged from 0.5 to 3.5), 16 participants in MP (scores ranged from 3.75 to 5), and 16 participants in HP (scores ranged from 5.25 to 6.5). As a sanity check, a one-way ANOVA on scores of the three groups showed a significant group effect (*F* = 84.32, *p* < 0.0001) and post-hoc Wilcoxon tests between pairs of groups indicated significant differences between HP and MP (*p* < 10^−6^) and between MP and LP (*p* < 10^−5^).

**Figure 1.**
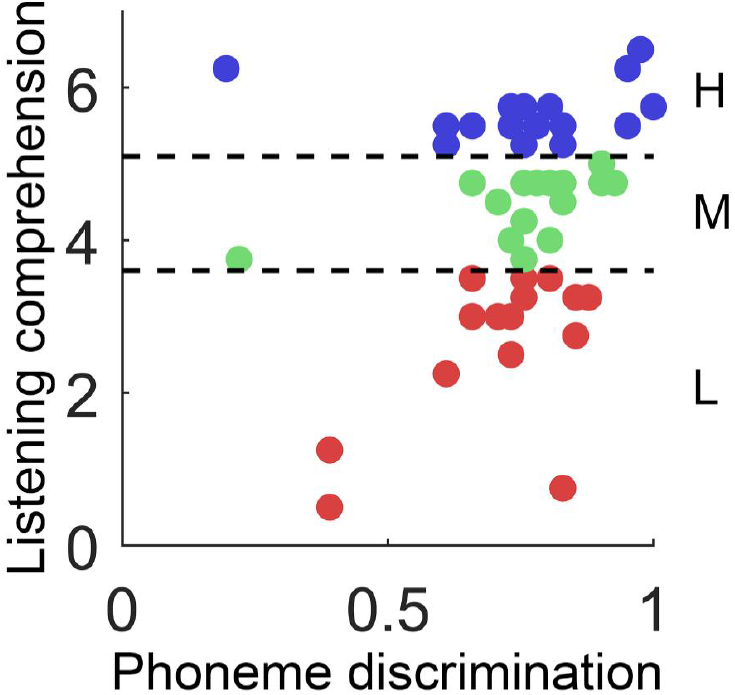
Scatterplot of scores on the Cambridge listening comprehension test and accuracy of phoneme discrimination for low, moderate and high proficiency groups. Dotted lines separate proficiency groups.

The mean phoneme discrimination accuracy of HP, MP and LP were 0.82 (SD = 0.12), 0.75 (SD = 0.16), and 0.71 (SD = 0.15). ANOVA revealed no significant effect of groups (*F* = 2.132, *p* = 0.131; Figure 1), suggesting similar lower-level abilities across groups.

Linear regressions using phoneme discrimination to predict comprehension scores showed that no significant relationship existed in HP or MP (*p*s > 0.05). In the LP group, the comprehension scores were found positively predicted by phoneme discrimination ability (*R^2^* = 0.29, *p* = 0.022), suggesting that overall language ability at the low end may depend on lower-level phonological processes.

### Neural signatures of second language proficiency

By using shared response model (SRM) in a searchlight procedure, we discovered neural synchronization across individuals at the same level of English proficiency during speech comprehension throughout the brain. For a given searchlight, if the response profile of one participant can be most accurately reconstructed based on the shared response of other participants at the same proficiency group, this region was considered characteristic to a proficiency level. Consistent responses for the high proficiency (HP) group were found throughout the bilateral medial and lateral frontal and occipital cortices, postcentral and precentral gyri, anterior cingulate cortices (ACC), posterior cingulate cortex/precuneus (PCC), insula, parahippocampal gyri (PHG), left intraparietal sulcus (IPS), and right supramarginal gyrus (SMG; Figure 2A). Compared with the moderate proficiency (MP) group, the HP group showed stronger synchronization in several clusters in the left anterior to middle temporal lobe (ATL), bilateral superior and middle frontal gyri, postcentral gyri, middle occipital gyri (MOG), right PHG, and right insula. Compared with the low proficiency (LP) group, the HP group showed stronger synchronization in bilateral lingual gyri, MOG, and right inferior frontal gyrus (IFG). In short, the HP group showed widespread synchronization in default mode network areas and in proposition representation areas.

**Figure 2.**
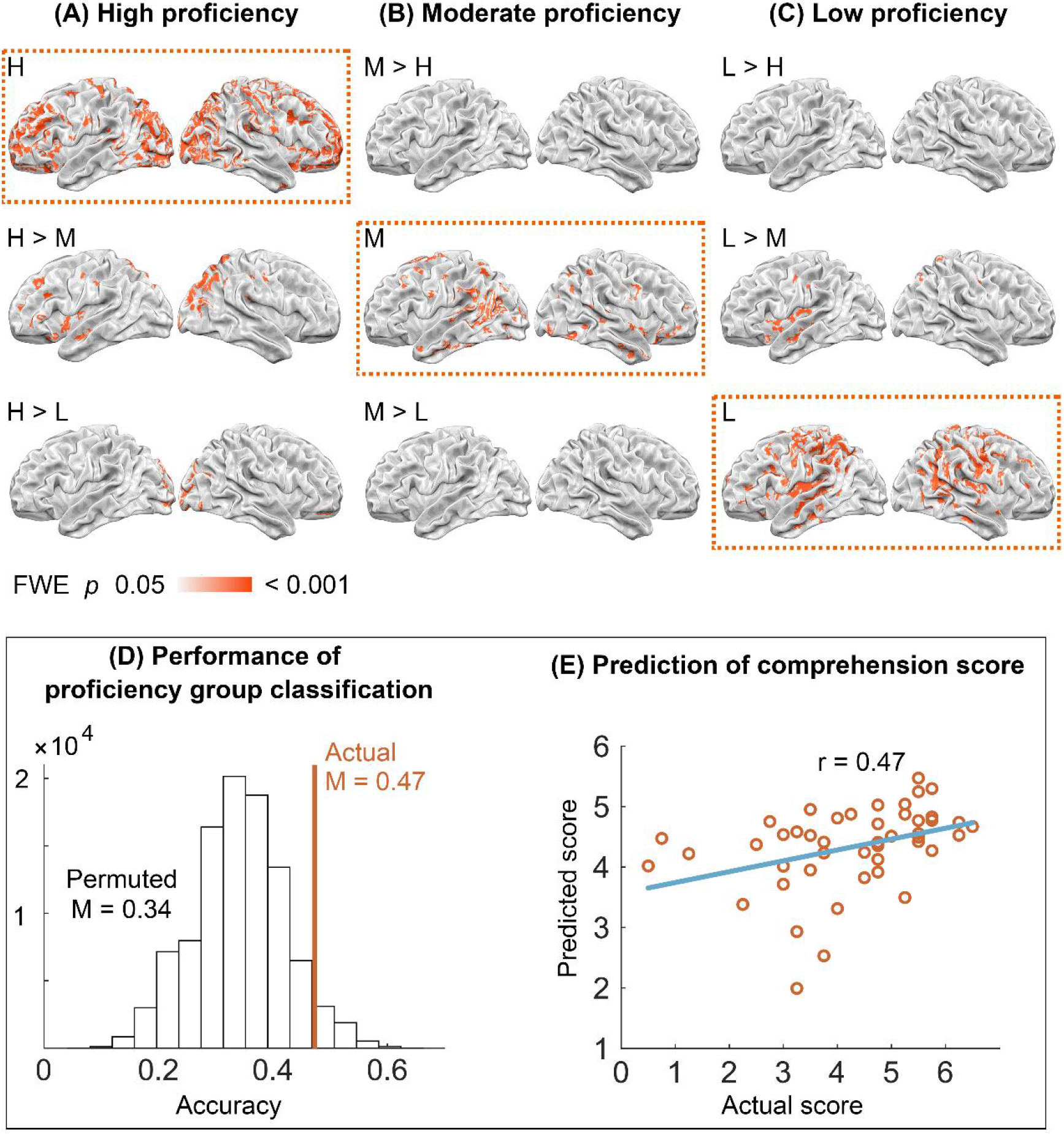
Proficiency-specific shared responses. Searchlight maps of neural synchronization across individuals for high **(A)**, moderate **(B)**, and low **(C)** proficiency groups. The three diagonal boxes show the proficiency-specific regions for each group, namely where a participant’s neural responses were most accurately reconstructed from the shared responses of other participants in the same group. The off-diagonal maps show where this same-group synchronization differed from the same-group synchronization of other groups. **(D)** Classification accuracy for predicting proficiency group from the synchronization maps of individual participants. The histogram shows the distribution of classification accuracies under null hypothesis of scrambling group membership. The vertical line shows the actual mean classification accuracy over participants. **(E)** Scatter plot of the actual and predicted scores in the Cambridge listening comprehension test.

For the MP group, consistent responses were found in the bilateral ATL, auditory cortices, posterior superior temporal sulci (pSTS), IPS, postcentral gyri, orbitofrontal cortex, ACC, PCC, PHG, left SMG, MOG, right IFG and lingual gyrus (Figure 2B). No regions showed greater synchronization for the MP group compared against either the HP or LP groups.

For the LP group, consistent responses were found in the bilateral auditory cortices, middle to posterior middle temporal gyri, IFG, precentral and postcentral gyri, ACC, PCC, superior parietal lobule (SPL), right insula, and lingual gyrus (Figure 2C). No regions showed greater synchronization compared with the HP group. The LP group showed greater synchronization in the left insula, Rolandic operculum, STS, bilateral precentral gyri, mPFC, PCC, right PHG and SPL.

### Predicting the proficiency level of individual participants

The mean accuracy of three-way classification that identified the proficiency group membership of individual participants was 0.49 (mean chance level at 0.33; *p* < 0.05; Figure 2D). The predicted English comprehension scores were also significantly correlated with the true comprehension scores over participants, *r* = 0.47, *p* = 0.008 (Figure 2E).

## Discussion

The present study found that inter-subject neural synchronization during second language narrative comprehension was broadly tuned by language proficiency. By using naturalistic continuous speech as the probe and searchlight SRM to align multi-subject responses with tolerance on local spatial idiosyncrasies, we identified proficiency-specific neural profiles that were more widely distributed than was previously thought.

The high proficiency group showed synchronization in regions that were similar to native speakers. Shared response profiles or activations across individuals during narrative processing in one’s native language have been repeatedly observed in the bilateral lateral frontal areas, mPFC, ACC, PCC, SMG, IPS, and ventral temporal areas (Chen et al., 2017; Honey et al., 2012; Lerner et al., 2011; Mazoyer et al., 1993; Nguyen et al., 2019; Simony et al., 2016; Yeshurun et al., 2017; Zadbood et al., 2017), many of which compose the default mode network (Raichle et al., 2001). The default mode network has been implicated in tracking information over longer timescales (Honey et al., 2012; Lerner et al., 2011) and representing high-level semantics (Chen et al., 2017; Nguyen et al., 2019; Yeshurun et al., 2017). A closer investigation has suggested that the PCC/precuneus is also engaged in higher-order information structures regardless of the semantic congruency, as long as the structure is consistent over time (Aly et al., 2018). Whereas synchronization in the PCC/precuneus was present in all groups, the proficient participants showed a particularly widespread synchronization in the medial and lateral frontal areas. The dorsomedial prefrontal cortex is one of the most robustly identified areas in various semantic processing tasks (Binder et al., 2009). Strong engagement of dorsomedial prefrontal cortex has been found in explicit phrasal composition (Graessner et al., 2021). The ventromedial prefrontal cortex is critical in integrating new information with prior schema (van Kesteren et al., 2010) and has been proposed to contribute to the very late stage of semantic composition that connects the core semantic systems with social cognition and episodic memory (Jackson et al., 2020; Pylkkänen, 2019). Thus, these results suggest that proficient L2 users rely on native-like neural systems during L2 speech processing, which incorporate regions beyond cognitive control areas or the core language network.

Previous studies have associated the improved L2 proficiency with reduced lateral prefrontal activations (Chee et al., 2001; Stein et al., 2009) and increased ACC-lateral frontal connectivity (Grant et al., 2015) during lexical-semantic processing. The present results showed that the activation profiles to continuous speech input in the dorsolateral prefrontal cortex, a typically observed area in selective attention and working memory tasks (Curtis and D’Esposito, 2003), were more alike among proficient individuals than the less proficient. These findings jointly suggest the speculation that for proficient L2 users, enhanced cognitive control from medial frontal cortex may increase local efficiency in lateral areas (decreased activation magnitude) and entrainment of lateral responses with the external narrative, increasing cross-participant synchronization.

The low proficiency group also showed considerable within-group consistency, mostly in the temporal and temporoparietal areas. The synchronized areas across these participants overlapped with phonological processing and the ventral lexical-semantic pathway proposed in the dual-stream model of speech perception (Hickok and Poeppel, 2007). Note that lower proficiency individuals showed statistically comparable ability in English phoneme discrimination as other groups, but only this group exhibited an association between behavioral performance on phonological and comprehension tasks. This suggests that participants with lower proficiency managed to perceptually track the continuous input and word-level semantics, but the narrative-level comprehension was disrupted or at least insufficient to reach agreement.

The moderate proficiency group showed the least internal consistency among the three groups, which coincides with findings that moderately proficient L2 can be associated with more deviations from the typical or L1-like activations (Dehaene et al., 1997; Mouthon et al., 2020). Interestingly, the low proficiency group showed greater synchronization than the moderate proficiency group in the temporal cortices as well as in the ACC and PCC. Considering that the participants were assigned to discrete groups based on their comprehension scores, originally a continuous measure, we speculate that the reduced consistency of moderate participants reflects greater heterogeneity in this group. If there is a state of ultimate attainment for L2 acquisition, a moderately proficient learner can be someone undergoing rapid development of skills, someone encountering a temporary bottleneck, or someone getting fossilized. Various types of intermediate states add to the diversity of the group compared to the experts or beginners.

A seeming surprising finding is the systematic response of the occipital lobe to speech processing. In fact, synchronized responses in lingual gyrus were seen in all proficiency groups. The involvement of lingual gyrus in sentence-level speech processing has been reported in various studies (Brennan et al., 2012; Hasson et al., 2018; Rodd et al., 2005; Zekveld et al., 2006). Indeed, there are several findings consistent with a role for occipital cortex in language proficiency. Congenitally blind individuals recruit occipital cortex in auditory sentence comprehension task (Bedny et al., 2011). Attention to speech input allows the content-specific responses to spread to the visual cortices (Regev et al., 2019). Higher-order correlations within visual areas are able to characterize content-specific inter-subject consistency during story listening (Owen et al., 2021). Cortical thickness of the lingual gyrus covaries with temporal cortex, inferior parietal cortex and inferior frontal gyrus (Chen et al., 2008). These findings are all consistent with the idea that early sensory cortices are modulated by or coactivate with higher-order areas, and may suggest that complex narrative processing is associated with information propagated across the brain.

Beyond descriptive findings, reliable accuracies in predicting the proficiency of individual participants confirmed that the neural signatures did reflect one’s L2 proficiency. Instead of using distilled and sometimes unnatural experimental comparisons based on postulated cognitive constructs, we examined neural response profiles in a naturalistic language comprehension task. This enabled a more detailed neural characterization at a finer-grained proficiency level. Moreover, using searchlight SRM allowed us to identify where in the brain there were consistent responses across participants without being restricted to a rigid voxel-to-voxel correspondence. This study has illustrated a promising application of the SRM technique, namely to identify individual differences via within-group commonalities and between-group differences in neural response profile.

## Acknowledgement

This work was funded by LAIX; the National Natural Science Foundation of China [31970987 to QC; 32100857 to JW]; the Basic Research Project of Shanghai Science and Technology Commission [No. 19JC1410101]; and the “Flower of Happiness” Fund Pilot Project of East China Normal University [No. 2019JK2203].

